# A miRNAs Based Exploration of promising Biomarkers in Cervical Cancer using Bioinformatic Methods

**DOI:** 10.1101/2021.12.27.474313

**Authors:** Elakkiya Elumalai, A. Malarvizhi, T. Sri Shyla, OM. Aruna devi, Krishna Kant Gupta

**Affiliations:** Centre for Bioinformatics, Pondicherry University, Pondicherry, India, 615014; School of Chemical and Biotechnology, SASTRA Deemed University, Tamil Nadu, India, 613401

**Keywords:** Cervical Cancer, Biomarker, miRNA, expression, pathways, protein-protein interaction

## Abstract

Cervical Cancer (CC) is a gynecologic cancer. In this cancer early detection is incredibly tough because most of the patients are not have any specific symptoms that results in suspending the proper identification. In this work, we selected TCGA CESC datasets and miRNA Seq analysis was done. The expression profiles of miRNAs in cervical cancer datasets were investigated using bioinformatics tools. The expression profiles of miRNA in Normal tissue, primary tumor and metastatic samples were analyzed. Based on p-value, principal component analysis and comparative literature survey, we reported 6 over-expressed (5X) miRNA at metastatic stage namely, hsa-mir-363, hsa-mir-429, hsa-mir-141, hsa-mir-93, hsa-mir-203b and hsa-mir-18a. Expression profiles were compared in heatmap. The target genes for the selected miRNAs were investigated for interaction and pathway details. The identification of two hub proteins (**PTEN and MYC**) in Protein-Protein Interaction Network was followed by pathway analysis. Our results indicate that **hsa-mir-363**, **hsa-mir-429, hsa-mir-141, hsa-mir-93, hsa-mir-203b and hsa-mir-18a** could be a potential diagnostic biomarkers for early-stage CESC and serve as prognostic predictors for patients with CESC.

## Introduction

Cervical Cancer is the highest common cancer that are faced by women all over the world. Early detection of cervical cancer is very difficult because most of the patients are not having any specific symptoms. Cervical cancer is the highest common gynecology cancer globally. It is a malignant tumor in cell of the cervix. Cancer in the cervix of the uterus is called cervical cancer. Cervix cell changes from normal to pre-cancer and then to cancer stage (Zhao *et al*. 2018) The primary underlying cause of cervical cancer is due to the infection of Human Papillomavirus, it is common virus called HPV which is transmitted during sexual activity. Human Papillomavirus (HPV) 16 and 18 have been found to cause 70% of cervical cancer causes. Most cases will be diagnosed in women between ages 35 and 44.The significant causes of cervical cancer is Human papilomavirus (HPV). This may take up to 20 years, or even longer days to develop cervical cells which are affected by HPV to cancerous tumor. Intake of vaccination against the most common HPV types associated with cervical cancer are the primary prevention. DNA testing and VIA (Visual Inspection With Acetic Acid) are alternative screening tests for cervical cancer prevention (Rath *et al*. 2016). Still, there is no promising biomarker for cervical cancer.

MicroRNAs represent a small non coding RNA which regulate messenger RNA for degradation and also for intercellular signaling. miRNAs act as a powerful biomarker for predicting responses and drug targets of cervical cancer (Kilic *et al*. 2015). It is an important role in gene expression and pathway regulation. miRNAs offers a great potential in medicine and gives treatment to various disease in future (Kori and Yalcin 2018). miRNAs and target genes can serve as biomarkers for cervical tumors which are associated with disease progression. Here miRNAs act as major role as it regulate gene expression as well as regulate biological process (Gao *et al*. 2018).

The miRNAs has a great potential in medicine and biomarker. In this present work, microRNA (miRNA) based biomarkers for early detection of cervical cancer were investigated. The 100 significantly differentially expresssed genes with a 85% variance in PCA1, were identified. The expression values of these differentially expressed genes were plotted in evolutionary heatmap. The target proteins of differentially expressed genes were identified, pathway enrichment analysis and protein-protein interaction was performed. After expression and pathway analysis, we proposed **hsa-mir-363**, **hsa-mir-429**, **hsa-mir-141**, **hsa-mir-93, hsa-mir-203b and hsa-mir-18a**, a promising biomarkers in cervical cancer.

### Clinical Significance

- We reported 6 over-expressed (5X) miRNA at metastatic stage as the important principal components namely, hsa-mir-363, hsa-mir-429, hsa-mir-141, hsa-mir-93, hsa-mir-203b and hsa-mir-18a.
- The main three pathways enriched for these six miRNAs were; pathways in cancer, hepatitis B and microRNAs in cancer
- We identified two hub proteins (**PTEN and MYC**) regulated by these ix miRNAs.

## 2. Methodology

### 2.1 Cervical Cancer datasets selection

Subio Plateform (Liu *et al*. 2013) was used to select datasets for cervical cancer and its expression analysis.

We took GDC miRNA Seq and we selected project “ TCGA CESC (Cervical Squamous Cell Carcinoma and Endocervical Adenocarcinoma)”. The workflow type was BCGSC miRNA. We collected 311 samples and successfully imported it in GDC miRNA-Seq plateform.

### 2.2 Data categorization and signal processing

We took median to calculate signals from same sample IDs. We created series and did normalization of count signals. The normalization includes filtering of signals whose count was less than 20 and global normalization with 95% percentile. We selected “cases.samples.sample_type” and added this column to our data table. It had three categories; a) Solid tissue normal b) Primary Tumor c) Metastatic. We set solid tissue normal as our control sample.

### 2.3 Signal filtering and differential gene expression analysis

We filtered signals in the range of −5 to 5. We filtered out those genes whose variance was less. The mi-RNA expression profiles of two samples; Primary Tumor and Metastatic was compared with control. The fold change was set to 5 and student’s T-test was used to compare the sample groups. The P-value was set to less than 0.05. Similarly, “Compare one to all” module was selected and upregulated-downregulated miRNAs were reported.

### 2.4 Heatmap and Principal component analysis

The datasets was analyzed for expression profiles of miRNAs in three different groups. The datasets was chosen to find the principal components (PC) contributing to the cervical cancer. We got majorly two principal components with their cumulative variance. Principal components Analysis (PCA) is a method for reducing the dimensionality without information loss (Jolliffe and Cadima. 2016).

### 2.5 Pathway enrichment analysis

Metascape server (Karnovsky *et al*. 2012) was used to perform Gene Ontology and KEGG pathway analysis of DEMs. Metascape is an analysis resource that helps to make sense of one or more gene lists. It provides automated meta-analysis tools for understanding either common or unique pathways and protein networks. A pathway has a set of genes related to a specific biological function and describes the relationship between the genes. This method helps for identifying biological pathways that are upgrade in a gene list that would be more than expected by chance.

### 2.6 Protein-Protein interaction analysis

The unique upregulated miRNAs in Normal Vs metastatic and Normal Vs Primary Tumor was considered and target proteins were selected from mirTarBase (Hsu *et al*. 2011). Protein-Protein interaction studies was done in STRING database (Szklarczyk *et al*. 2017). This database helps for analyzing familiar protein-protein interactions. The output was the Protein-Protein interaction (PPI) Network in the .tsv file format. Pathway enrichment analysis of Normal Vs metastatic and Normal Vs Primary Tumor was explored in STRING database.

## 3. Results and Discussion

### 3.1 Cervical Cancer Dataset selection, normalization and filtering

The “**TCGA CESC (Cervical Squamous Cell Carcinoma and Endocervical Adenocarcinoma)**” was having 311 samples. The samples were imported and the miRNA Seq count was normalized (Figure 1). There were 3 “Solid tissue Normal”, 306 “Primary Tumor” and 2 “Metastatic”. The three groups were separated and colored differently (Figure 2).The miRNAs whose count was less than 10 was filtered out (Figure 3). Out of 1881 miRNAs, only 382 miRNAs passed this filter (Figure 3).

### 3.2 Differential miRNAs expression across all the three groups

The “Solid tissue Normal” was selected as control and compared with “Primary Tumor” and “Metastatic” (Figure 4). The “Solid tissue Normal” was taken as reference and compared with “Primary Tumor” and “Metastatic”. At first, 5X upregulation with p-value < 0.05 analysis was done. There were 29 miRNAs showing upregulation as compared with Primary Tumor. Similarly, there were 17 miRNAs showing upregulation as compared with metastatic. Thereafter, 5X downregulation with p-value < 0.05 analysis was done. There were 16 miRNAs showing downregulation as compared with Primary Tumor. Similarly, there were 15 miRNAs showing downregulation as compared with metastatic (Figure 5). The names of miRNAs are given in Table 1.

### 3.3 Unique and common miRNAs in downregulated and upregulated datasets

We compared downregulated miRNAs in Primary tumors and metastatic with respect to solid normal tissue. There were 3 unique miRNAs in Normal Vs metastatic namely hsa-mir-29a, hsa-mir-1247 and hsa-mir-582. The hsa-mir-29a is shown to inhibits the metastasis and invasion of cervical cancer (Gong *et al*. 2019). The has-mir-1247 is shown to inhibits cell proliferation by targeting neuropilins (Shi e*t al*. 2014). The involvement of hsa-mir-582 in cervical cancer is reported by Chen *et al*. 2018.

There were 4 unique miRNAs in normal vs primary tumor namely hsa-mir-140, hsa-mir-381, hsa-mir-139, hsa-mir-204. The hsa-mir-140 inhibits the proliferation of human cervical cancer by targeting RRM2 (Ma *et al*. 2020). The hsa-mir-381 regulates the invasion of human cervical cancer cells by targeting G Protein Coupled Receptor 34 (GPR 34) (Tan *et al*. 2021). The decreased expression of hsa-mir-139 was found in cervical cancer (Sannigrahi *et al*. 2017). In lung cancer, the decreased expression of hsa-mir-204 was reported (Liang *et al*. 2020).

There were 12 common miRNAs in both datasets namely, hsa-mir-10b, hsa-let-7c, hsa-mir-1-2, hsa-mir-143, hsa-mir-99a, hsa-mir-100, hsa-mir-145, hsa-mir-133a-1, hsa-mir-133a-2, hsa-mir-1-1, hsa-mir-125b-1 and hsa-mir-125b-2. The downregulation of hsa-mir-10b is reported in small cell carcinoma of cervix by Huang et al., (Huang *et al*. 2012). In general, people proposed hsa-let-7c as a promising biomarker in cancer (Chirshev *et al*. 2019). Noone has reported the association between cervical cancer and hsa-mir-1-2, hsa-mir-1-1. The hsa-mir-143 act as cervical cancer suppressor gene (Liu *et al*. 2012). Noone has validated the downregulation of hsa-mir-99a in cervical cancer. Decreased expression of hsa-mir-100 was found in cervical cancer (Li *et al*. 2015). The downregulation of hsa-mir-145 have been reported in cervical cancer (Ma and Li 2019). The hsa-mir-133a targets EGFR and inhibits cervical cancer growth (Song *et al*. 2015). The expression level of hsa-mir-125b was altered in HPV Infection and Cervical Cancer Development (Ribeiro et al. 2015).

Similarly, We compared upregulated miRNAs in Primary tumors and metastatic with respect to solid normal tissue. There were 4 unique miRNAs in Normal Vs metastatic namely hsa-mir-34c, hsa-mir-93,hsa-mir-106b and hsa-mir-18a. In our study, we found, the overexpression (5X) of hsa-mir34c in metastatic cervical cancer cell but in contrary Sommerova et al., reported the underexpression of hsa-mir34c in cervical cancer cells (Sommerová *et al* 2018). The upregulation of hsa-mir-106b have been reported in cervical cancer (Yi *et al*. 2018). The suppression of hsa-mir-93 inhibits HPV positive cancer cell progression (Li *et al*. 2019), in our study it was found to be overexpressed (5X) and helps in cervical cancer progression. It could be a promising biomarker in cervical cancer. The dual role of hsa-mir-18a in promoting cancer or inhibiting cancer have been reported (Shen *et al*. 2019) but no one specifically reported its overexpression and exact role in cervical cancer.

There were 16 unique miRNAs in normal vs primary tumor namely hsa-mir-142,hsa-mir-200b, hsa-mir-944,hsa-mir-16-2,hsa-mir-15b,hsa-mir-425, hsa-mir-16-1, hsa-mir-155, hsa-mir-32,hsa-mir-200a, hsa-mir-135b, hsa-mir-224, hsa-mir-203b, hsa-mir-196a-2, hsa-mir-1307 and hsa-mir-200c. The lower expression of hsa-mir-142 was found in cervical cancer tissue (Li *et al*. 2019) but in our study it was found to be overexpressed (5X). The silencing of hsa-mir-200b reduced the growth of cervical cancer tissue (Wang and Chen 2019). In our study it was found to be overexpressed. The hsa-mir-944 has been reported as a biomarker poor prognosis of advanced cervical cancer (Park *et al*. 2019).Noone has reported the specific association between hsa-mir16-2, hsa-mir16-1 and cervical cancer. The hsa-mir-15b is associated with cervical cancer. The hsa-mir-425 is upregulated in renal cancer (Quan *et al*. 2018) but no reports are there for its association with cervical cancer. The overexpression of hsa-mir-155 is associated with increased risk of cervical cancer in HPV E6/E7 mRNA positive tissues (Park *et al*. 2017). The hsa-mir-32 was reported to be downregulated in cervical cancer but in our study it is upregulated (Liu *et al*., 2019). The overexpression of hsa-mir-135b has been reported in oral and lung cancer (Lopes *et al*. 2018). The hsa-mir-224 inhibits autophage and promotes cervical cancer (Fang *et al*. 2016).In our study, we found the overexpression of hsa-mir-203b. In cervical cancer, miR-196a inhibits p27kip1, FOXO1 and promotes cell proliferation (Lu *et al*. 2016). The upregulation of hsa-mir-1307 has been reported in breast and ovarian cancer by targeting SMYD4 protein (Han *et al*. 2019).

There were 13 common miRNAs in both datasets namely hsa-mir-183, hsa-mir-203a, hsa-mir-20b, hsa-mir-31, hsa-mir-182, hsa-mir-96, hsa-mir-141, hsa-mir-130b, hsa-mir-429, hsa-mir-106a, hsa-mir-210, hsa-mir-363 and hsa-mir-205. The hsa-mir-183 is associated with several cancer (Cao *et al*. 2020). The hsa-mir-203a is not specifically associated with cervical cancer. In High Grade Cervical Intraepithelial Neoplasia, the expression of hsa-mir-20b was found to be high (Szekerczés *et al*. 2020). The hsa-mir-31 was found to be upregulated in cervical cancer (Wang *et al*. 2017). The hsa-mir-182 plays an onco-miRNA role in cervical cancer (Tang *et al*. 2013). The hsa-mir-96 enhances tumorigenicity of human cervical carcinoma cells through PTPN9 (Ma *et al*. 2018).The hsa-mir-141 inhibits colorectal cancer by targeting TRAF5 (Liang et al. 2019) but we found in our study, it is overexpressed in cervical cancer. It may be playing protective role. The hsa-mir-130b targets TNF-α and promotes carcinogenesis of cervical cancer (Zhang *et al*. 2014).The hsa-mir-429 inhibits CDKN2B and promotes bladder cancer (Yang *et al*. 2017) but its overexpressed status in cervical cancer is unknown. The hsa-mir-210 was upregulated in cervical cancer. It was proposed to be used as micro RNA signature for cervical cancer detection (Liu *et al*. 2018). The hsa-mir-363 was found to exhibit protective role in ovarian cancer as its overexpression decreased growth, colony formation, migration and invasiveness of SKOV3 cells (Lin *et al*. 2017). Therefore, in cervical cancer also it might be playing protective role. The serum hsa-mir-205 was reported as novel biomarker for cervical cancer patients (Ma *et al*. 2014).

### 3.4 Heatmap and Principal Component analysis

A Heatmap is represented in form of graphical data that uses color coding to represent values. Heatmap shows the relative intensity of expression values. Variation of colors depends on its intensity value. Here red indicates over expressed regions, grey represents less expressed regions and whereas blue denotes normal expressed regions. Average linkage algorithm helps to group the distance between the weighted values so that two groups have an equal influence on result part. Pearson correlation method (Zhao *et al*. 2014) is utilized to see the linear relationship between the two quantitative variables. Finally, heatmap is displayed and also along with heatmap row dendogram in form of tree structure is also designed (Figure 6). In order to identify significant miRNAs, Principal component analysis was done. PC1 contributed to the variance of 85.52% and PC2 contributed to the cumulative variance of 14.47% (Figure 7). The 100 important miRNAs in PCA1 are shown in Table 2 & Figure 8.

### 3.5 Gene annotation description analysis

The 100 miRNAs from PC1 were selected for pathway enrichment analysis and analyzed the connected biological pathways and Gene Ontological annotations. The enriched term for 100 genes are given in Figure 9. The process enrichment analysis was given in Table 3. The main enriched terms were microRNAs in cancer, miRNA involved in DNA damage response, regulation of angiogenesis, regulation of STAT cascade and negative regulation of cell migration.

### 3.6 Target genes collection and pathway analysis

The 14 experimentally validated with strong evidence, target genes for over-expressed miRNAs in Normal Vs metastatic were collected. The protein-protein interaction of these proteins are shown in Figure 10. The main three pathways enriched were; pathways in cancer, hepatitis B and microRNAs in cancer (Table 4). Similarly, 27 target genes for over-expressed miRNAs in Normal Vs primary tumor were collected. The protein-protein interaction of these proteins are shown in Figure 11. The main three pathways enriched were; pathways in cancer, proteoglycans in cancer and microRNAs in cancer (Table 5).

## 4. Conclusion

We have used different subio plateform to analyze differential miRNA gene expression values in cervical cancer datasets which was having 311 samples. We normalized the count with global normalization of 95%. We filtered out those signal whose count was less than 10. We set t-test, p value < 0.05 and fold change of 5. Finally, we has 381 significant miRNAs. We performed differtial gene expression analysis in primary tumor and metastatic with reference to the normal tissue. There were 29 miRNAs showing upregulation as compared with Primary Tumor. Similarly, there were 17 miRNAs showing upregulation as compared with metastatic. Similarly, there were 16 miRNAs showing downregulation as compared with Primary Tumor. Similarly, there were 15 miRNAs showing downregulation as compared with metastatic. The main enriched GO terms were microRNAs in cancer, miRNA involved in DNA damage response. Based on expression, pathway, principal component analysis and literature survey, this study reported 6 over-expressed (5X) miRNA at metastatic stage namely, hsa-mir-363, hsa-mir-429, hsa-mir-141, hsa-mir-93, hsa-mir-203b and hsa-mir-18a. It could be used as appropriate biomarker for the earlier detection of Cervical cancer but its clinical validation is required.

## Acknowledgment

We sincerely acknowledge SASTRA Deemed University for providing computational resources.

## Ethics with regard to experiments

No animals or living organisms were used in this study.

## Data Availability Statement

Data will be made available upon request to the corresponding author.

---

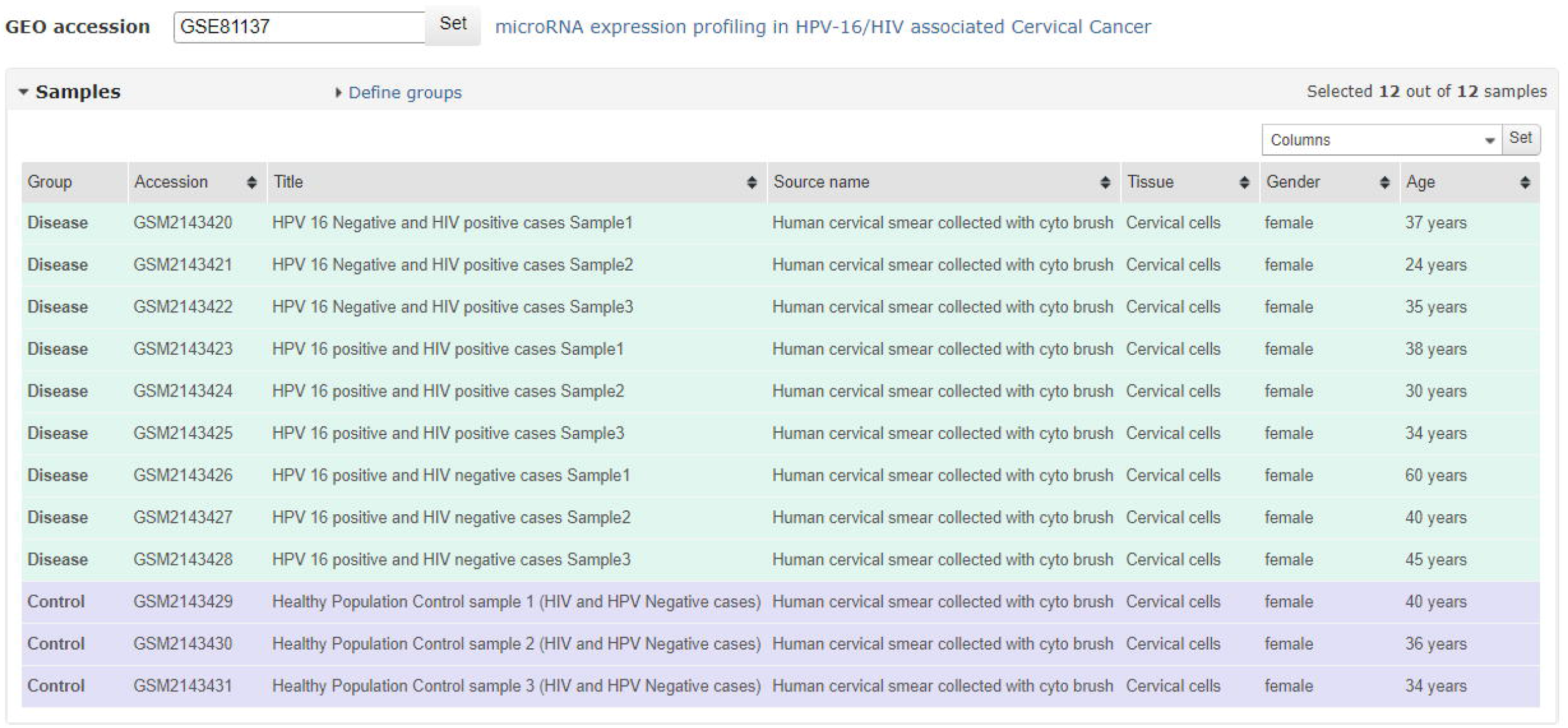

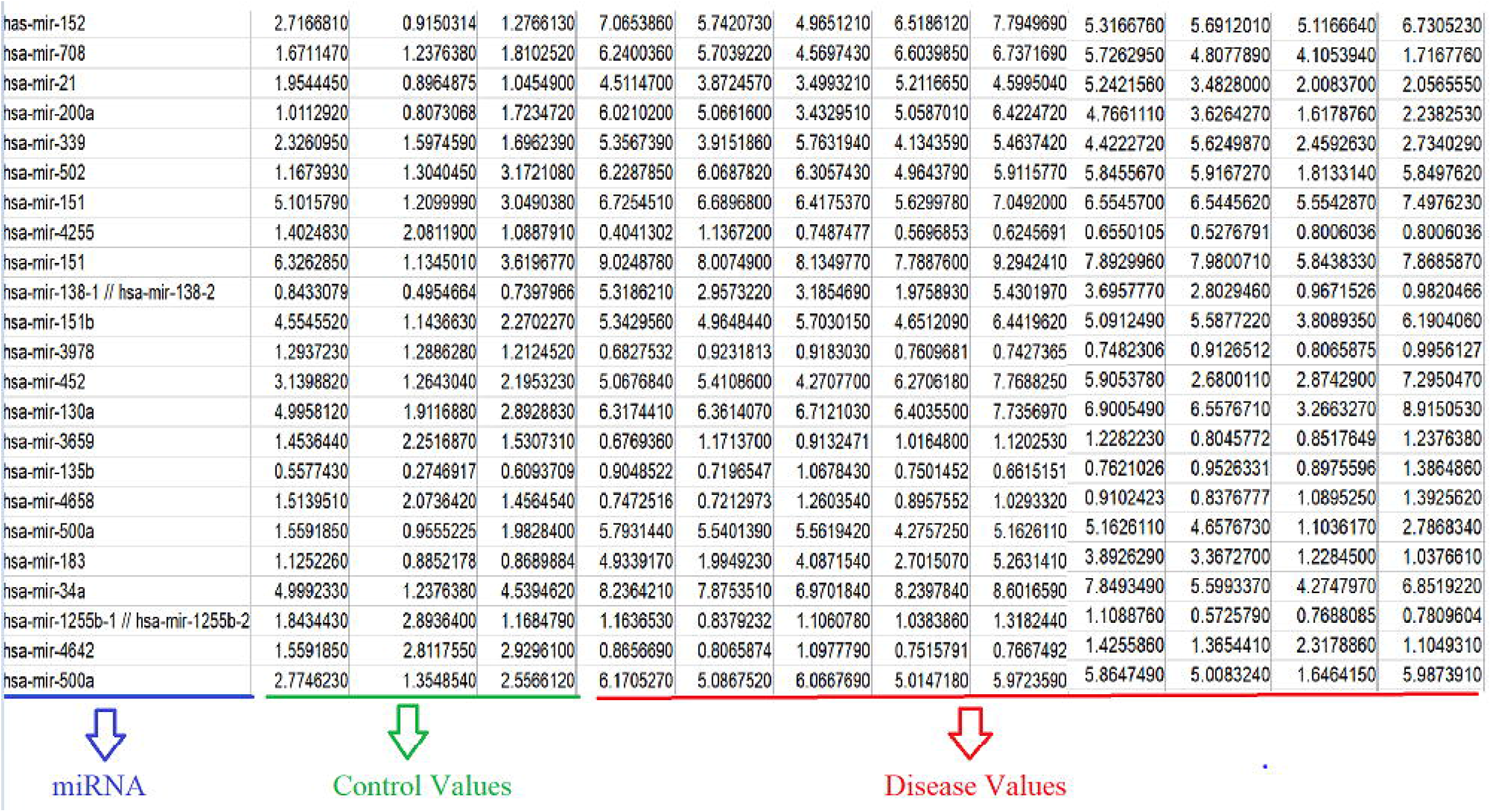

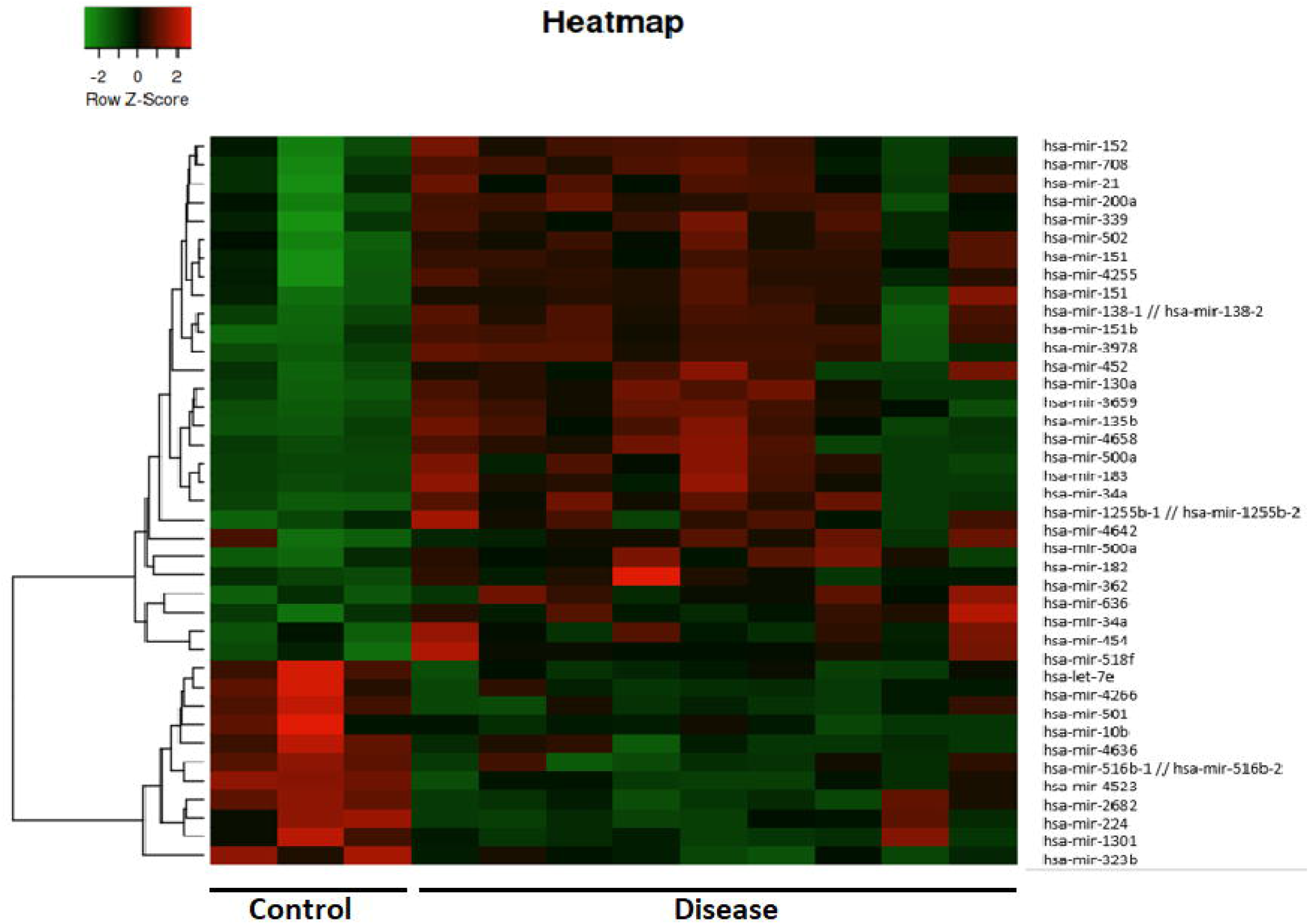

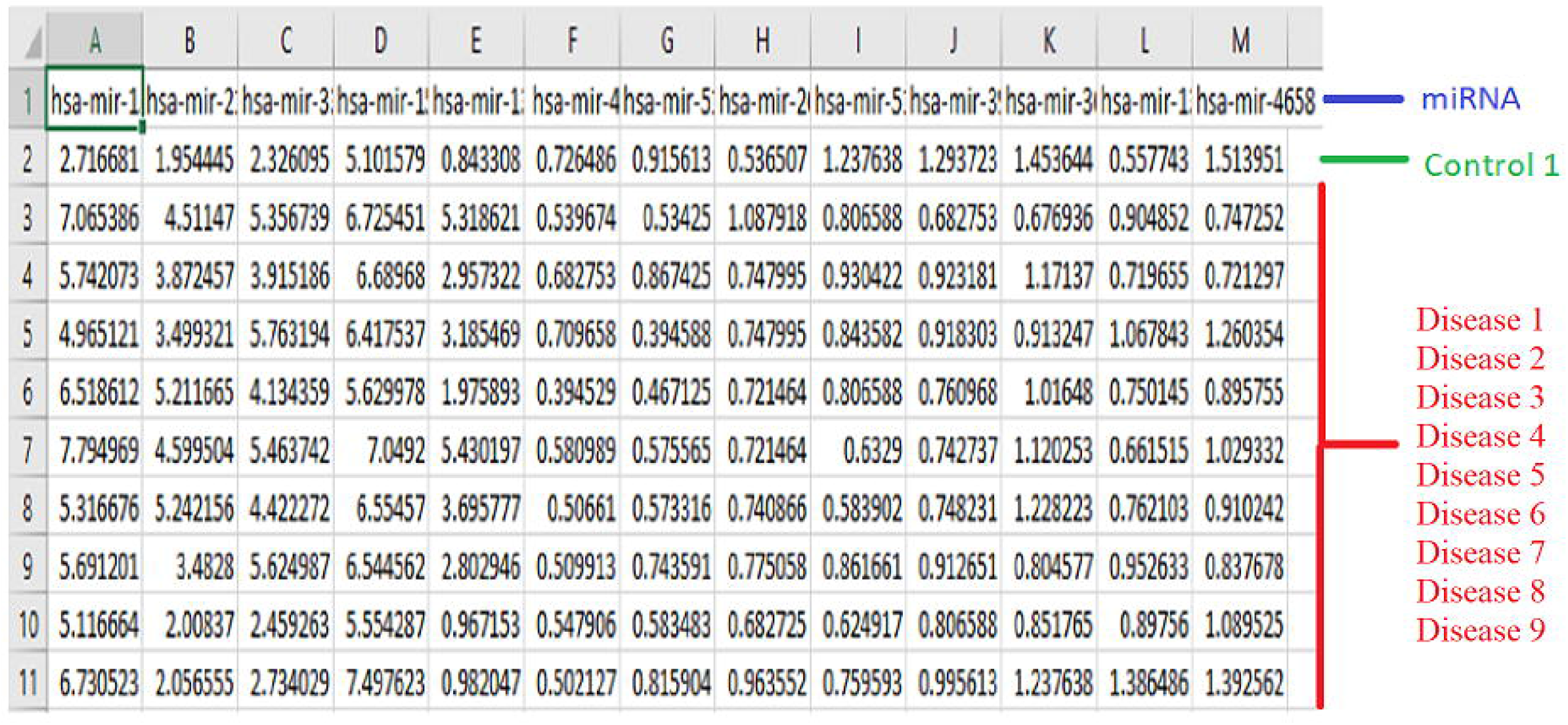

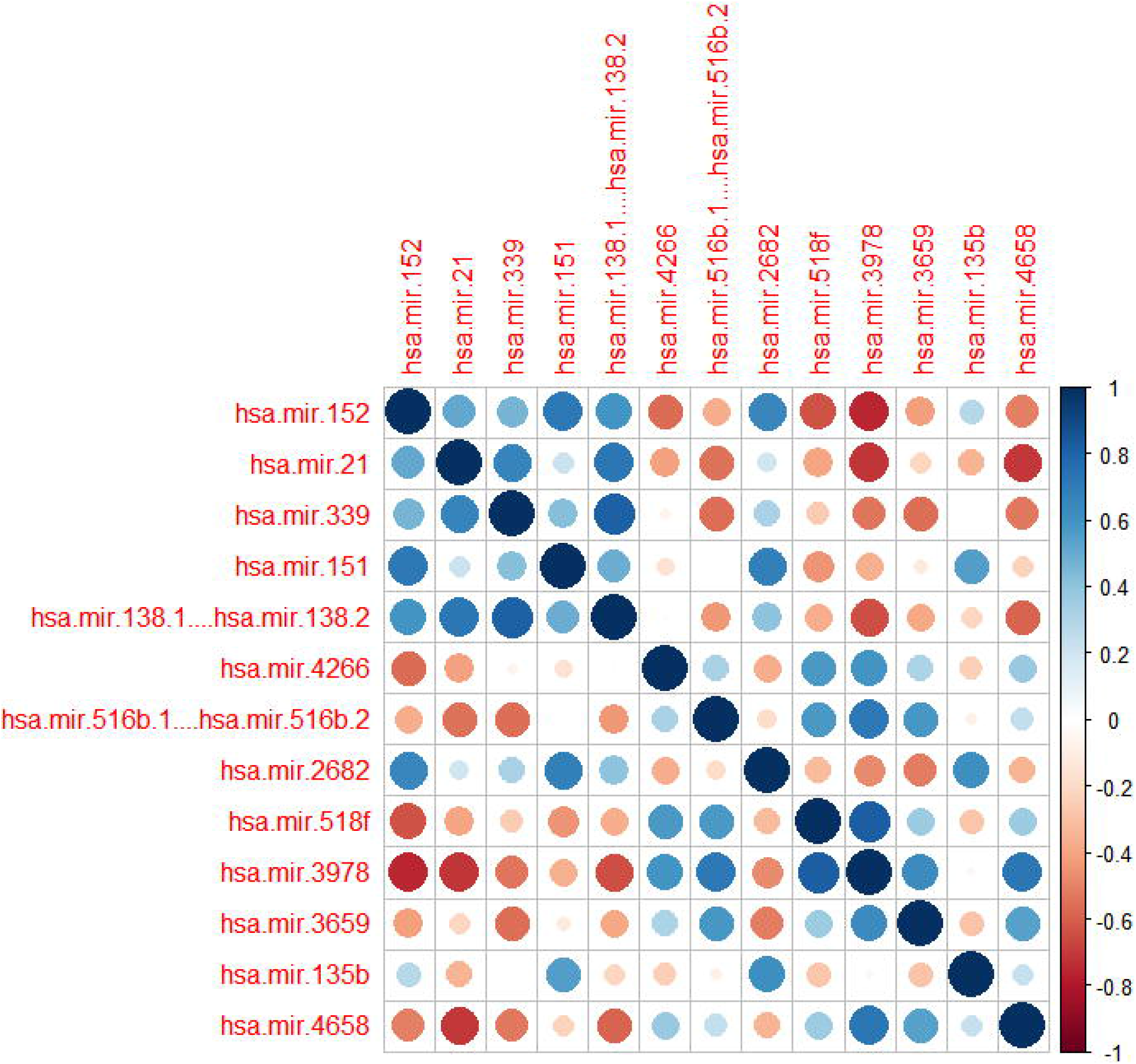

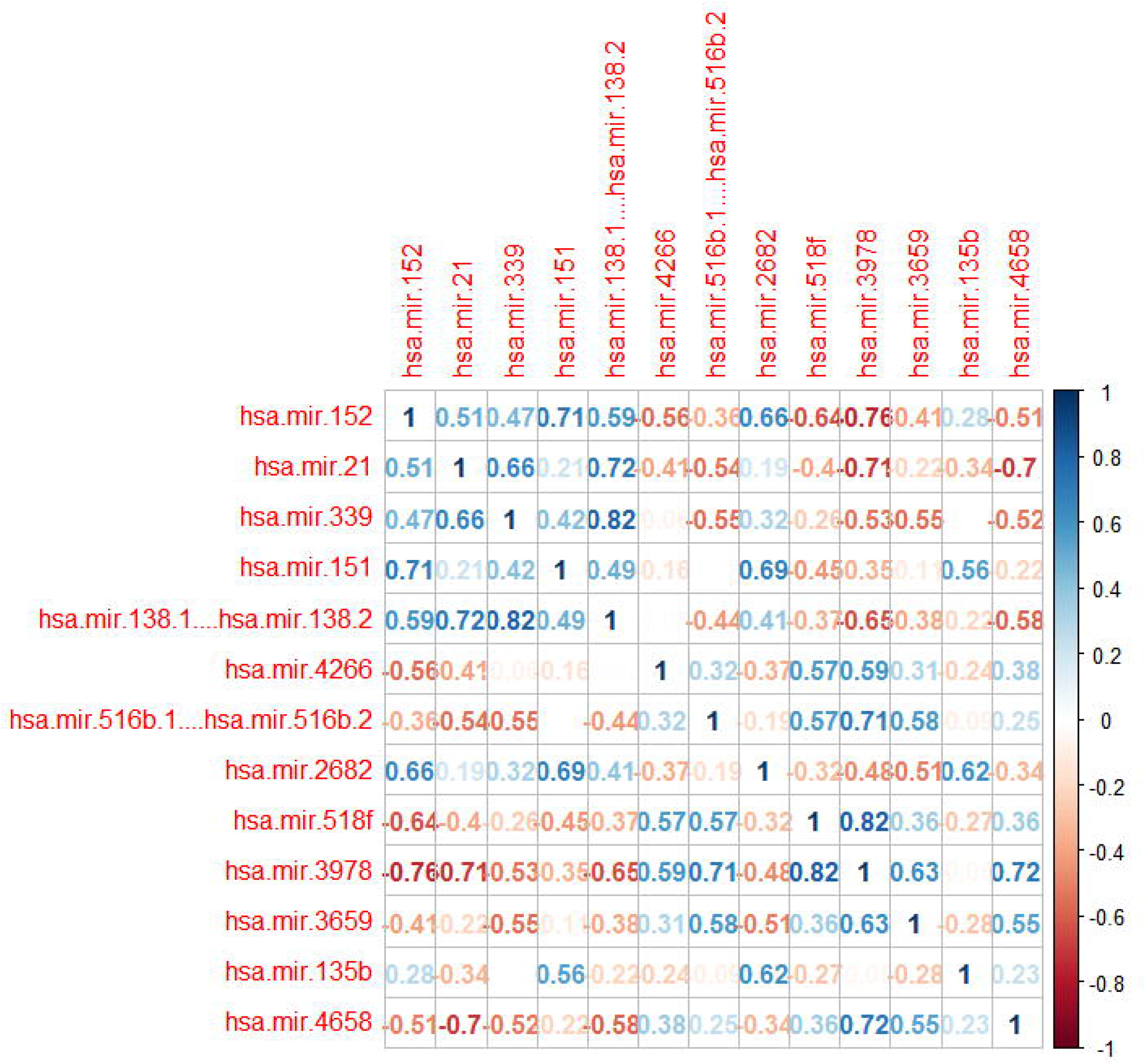

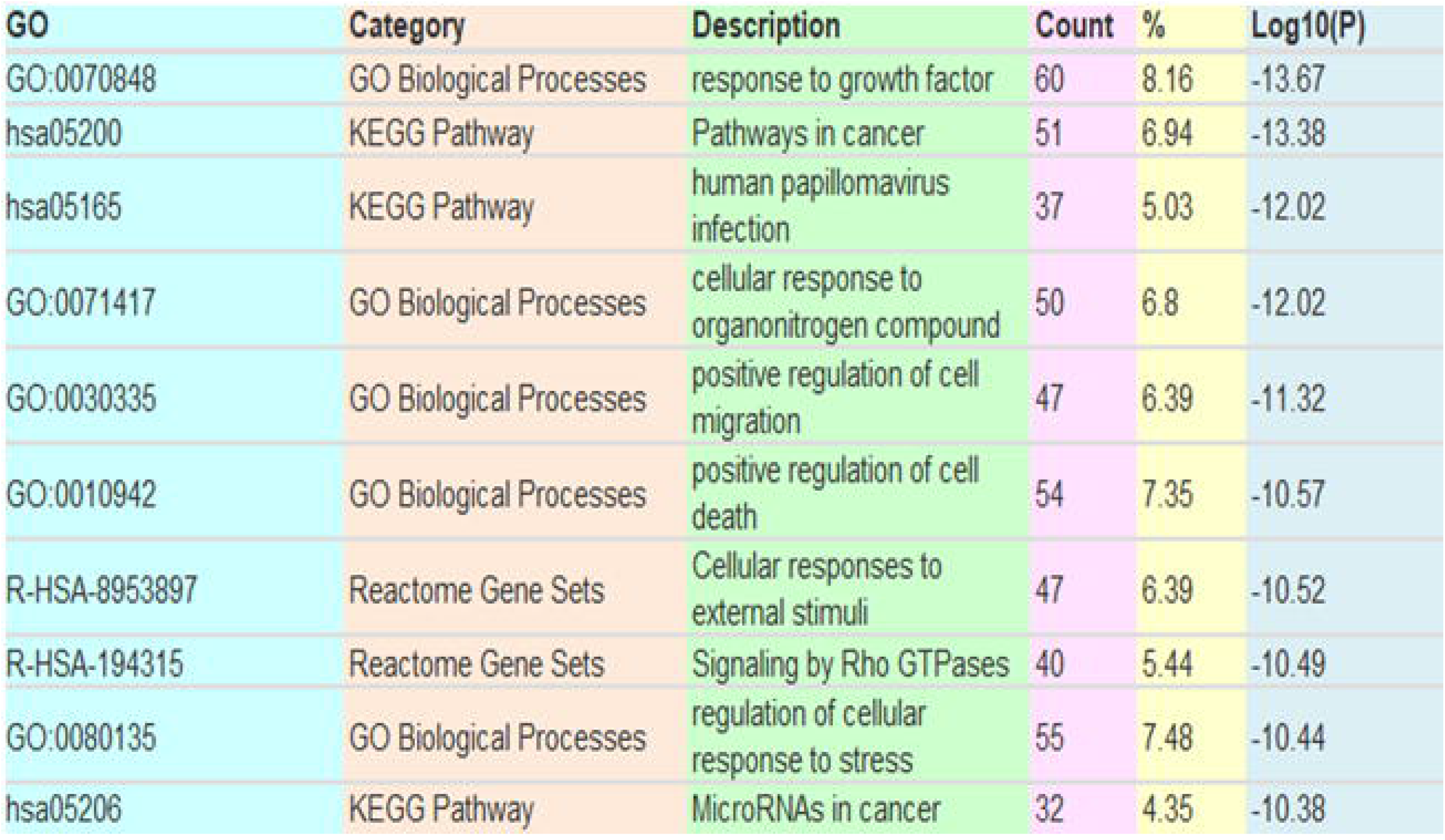

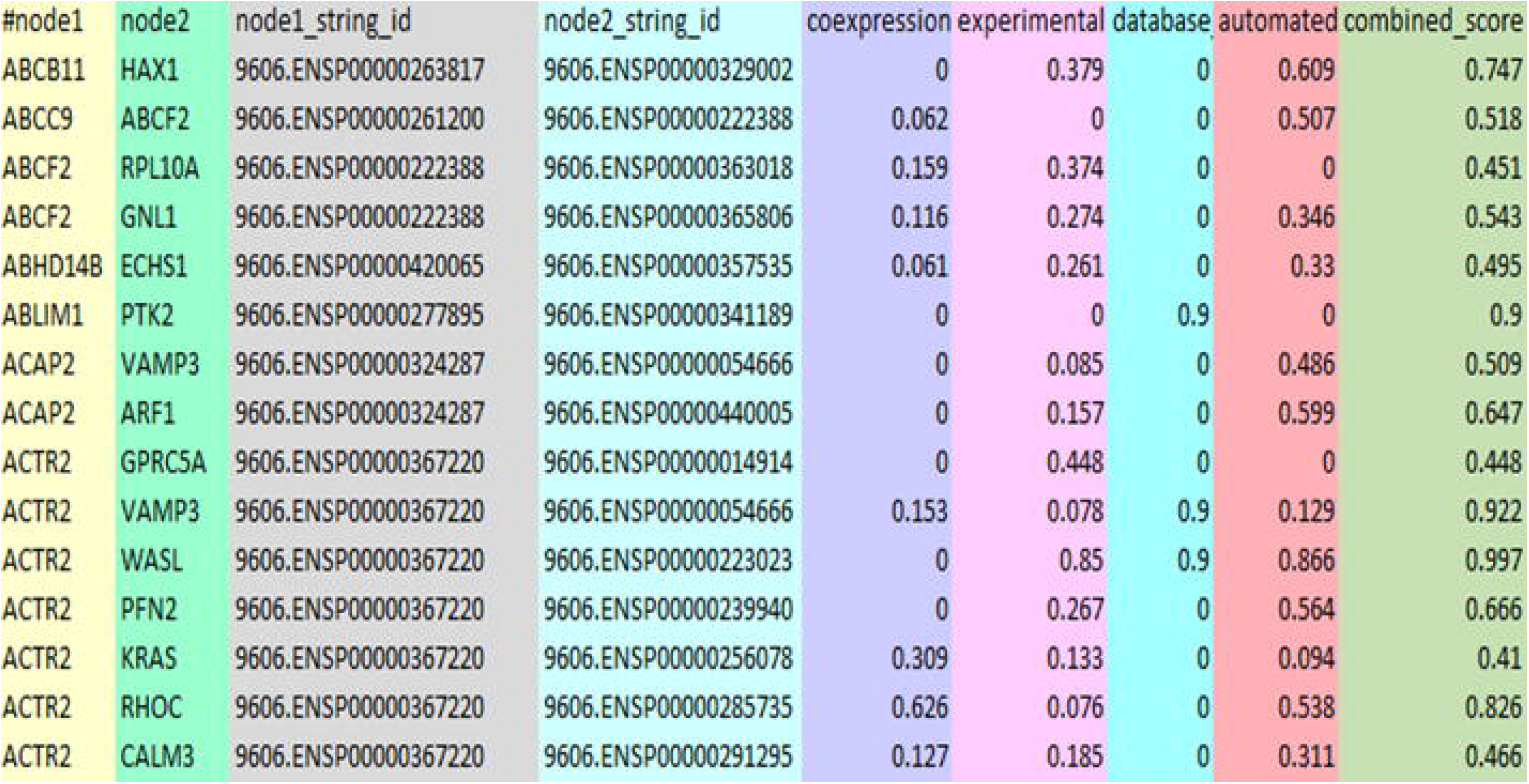

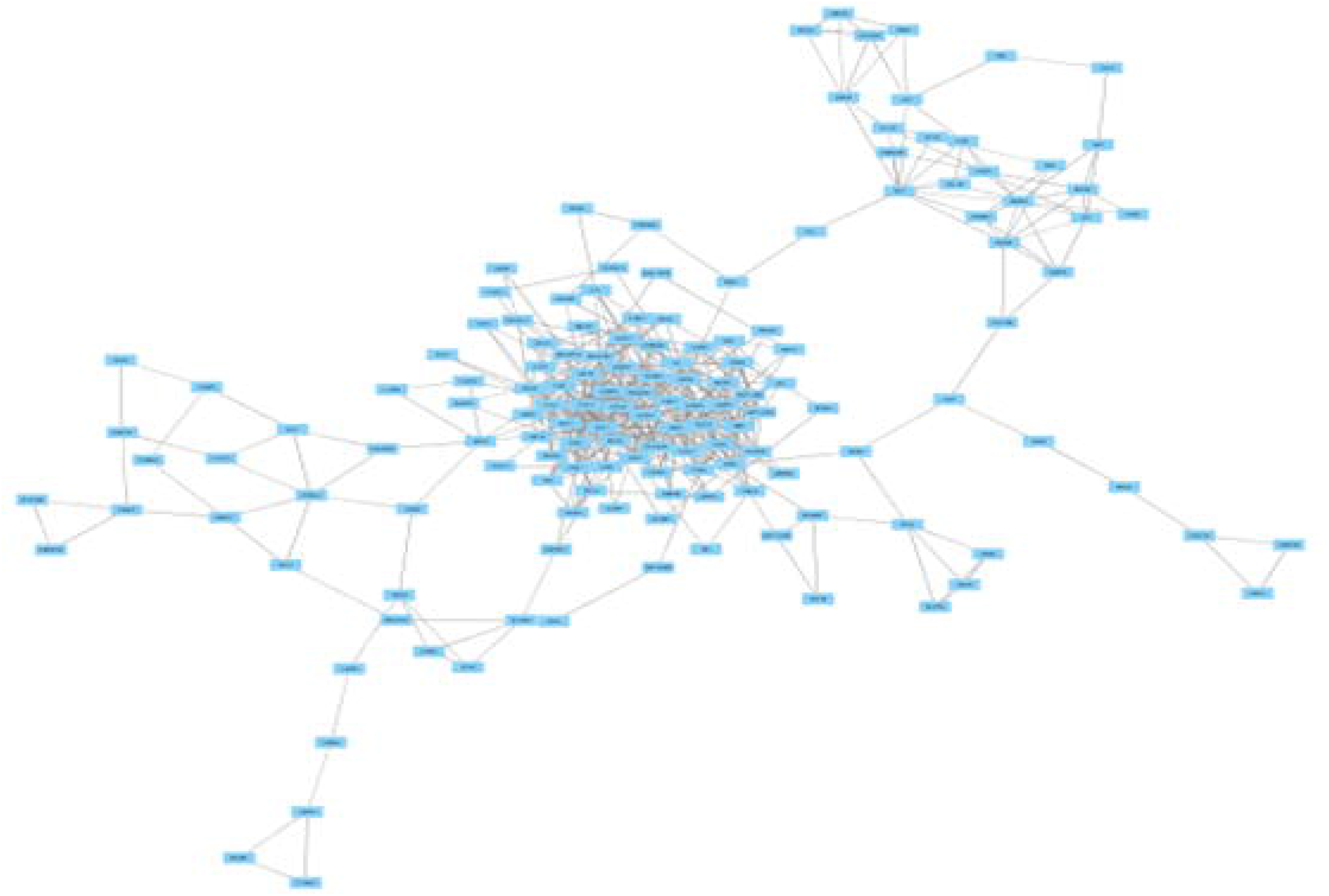

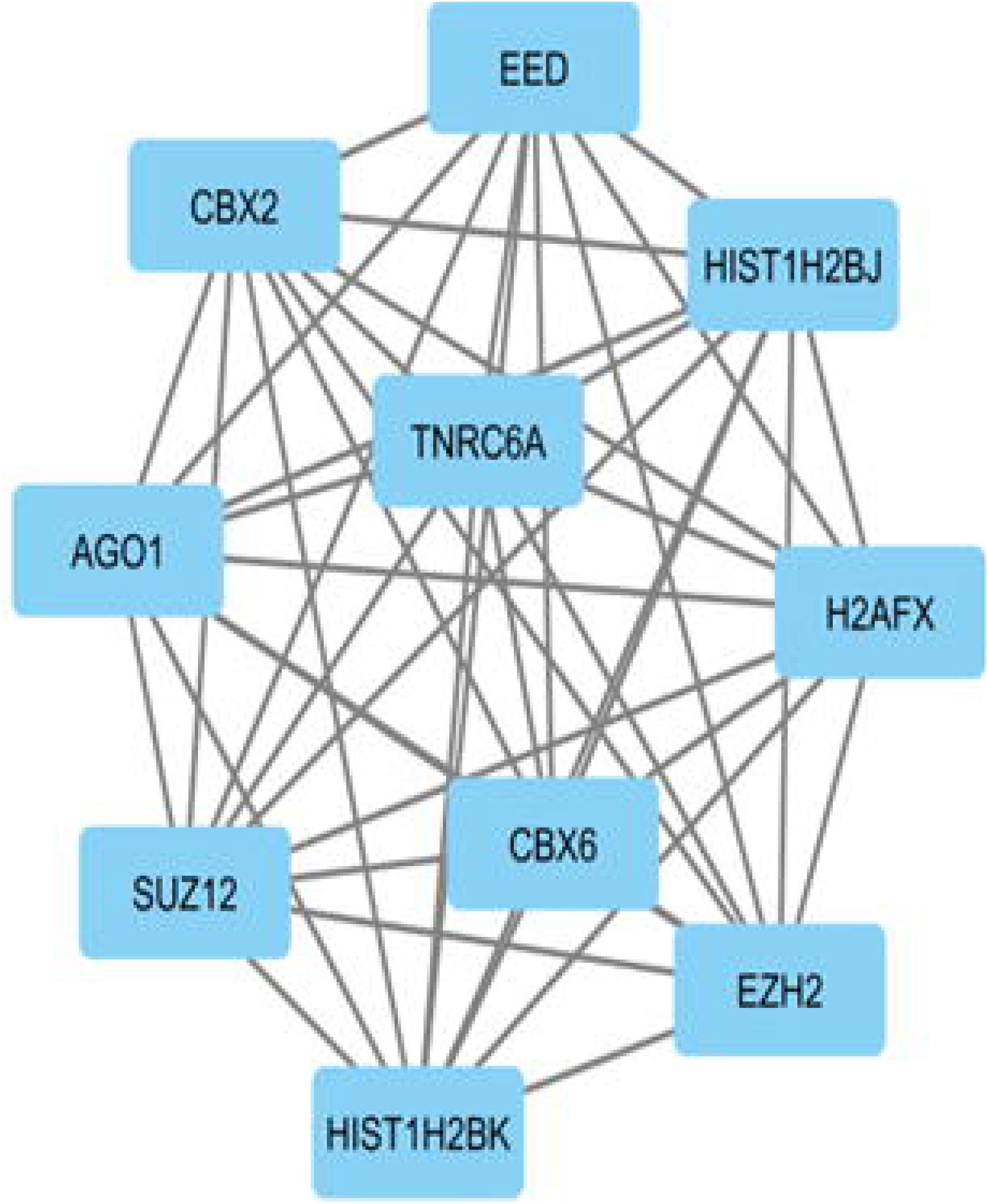

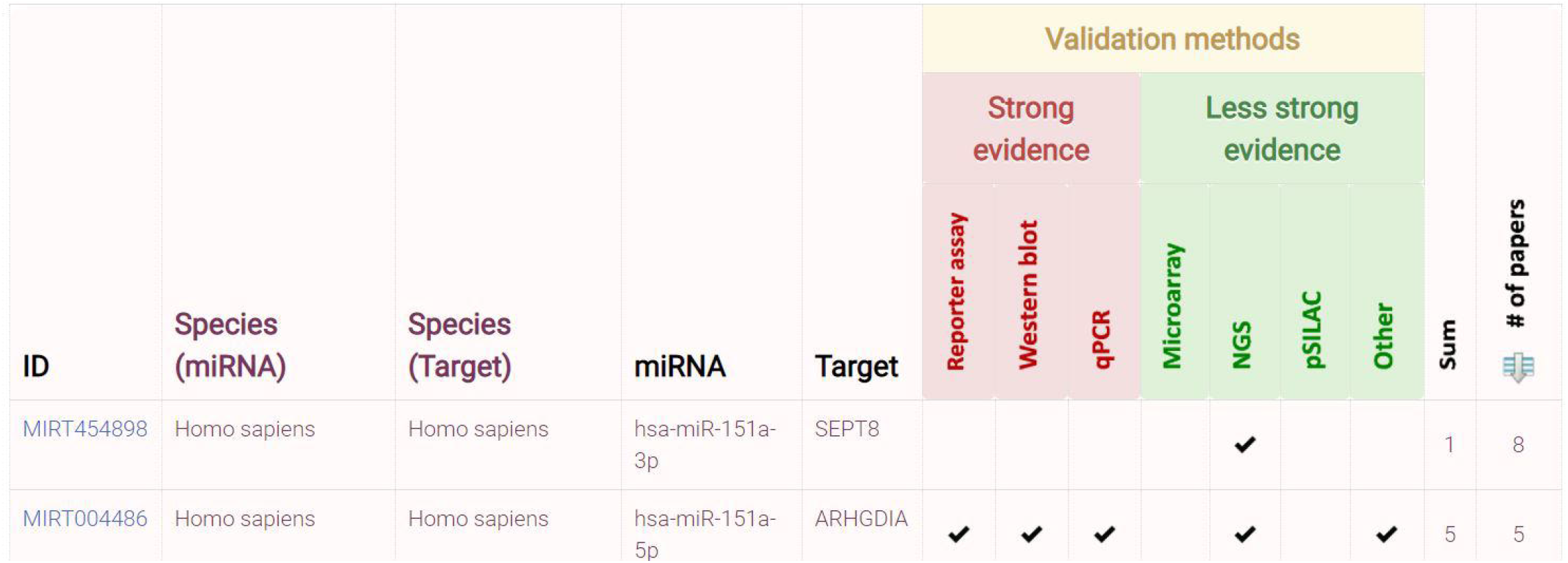

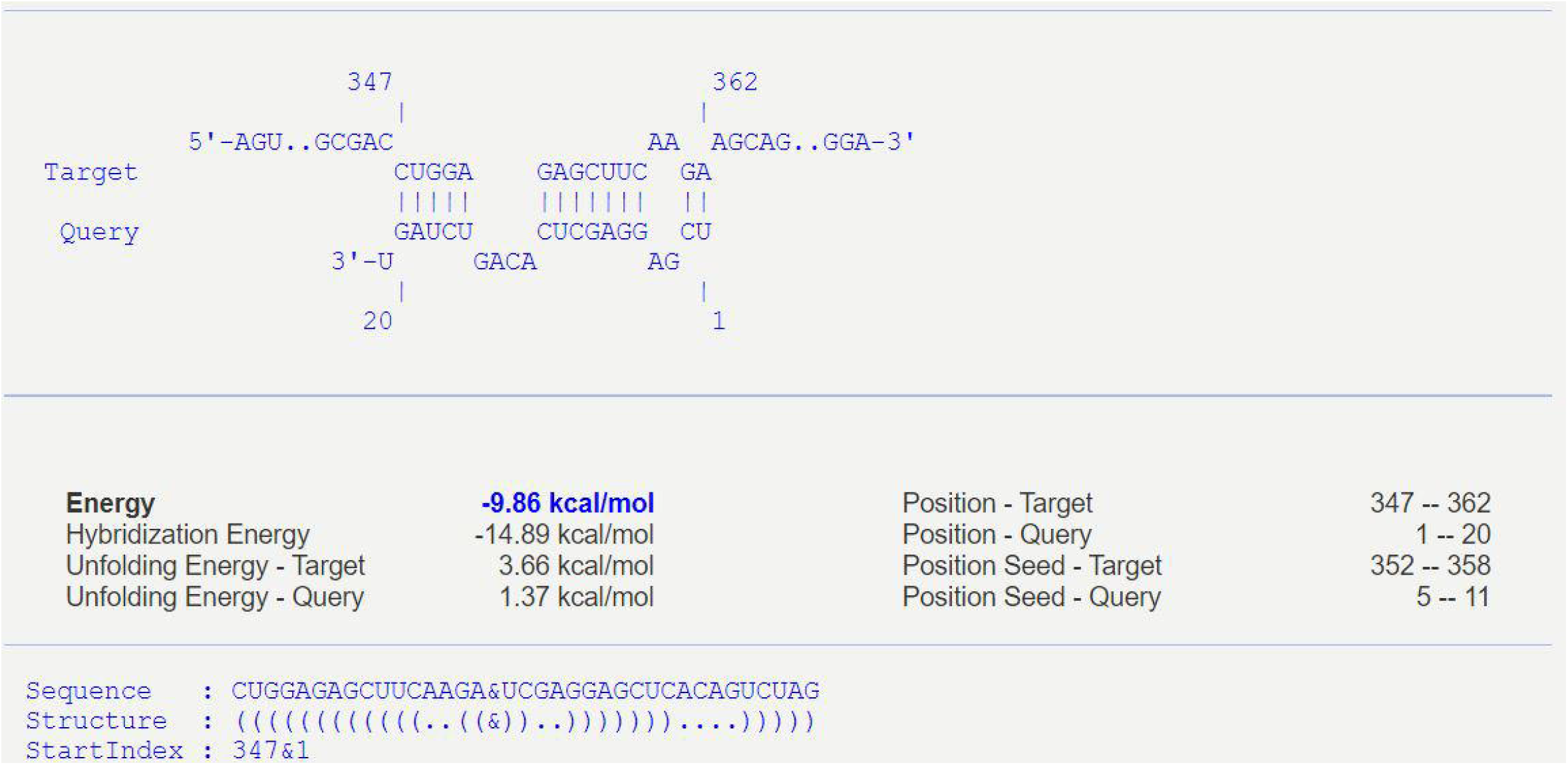

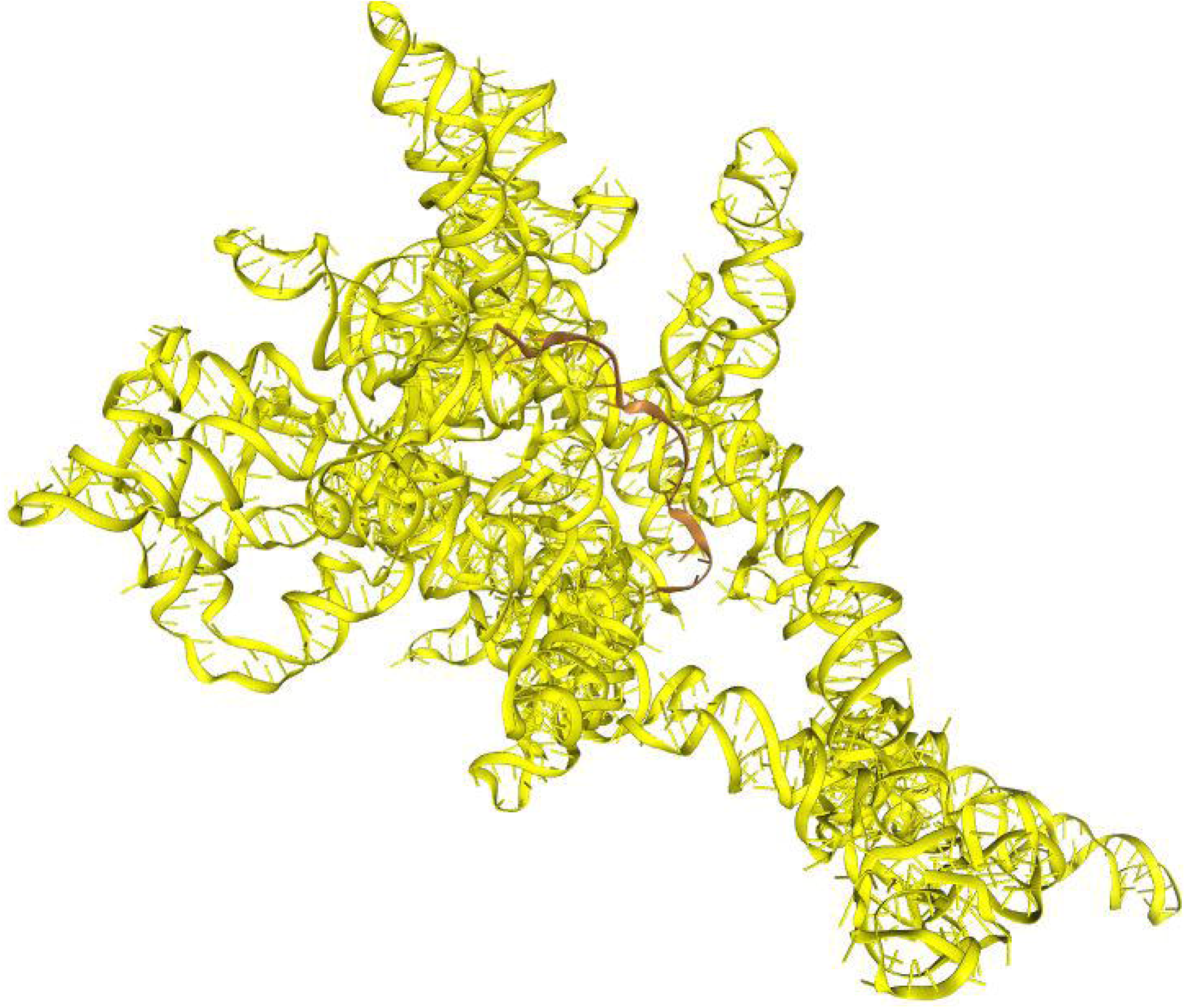

