## Supplementary material for "A miRNAs Based Exploration of promising Biomarkers in Cervical Cancer using Bioinformatic Methods": Table 1

| **miRNA_ID_LIST** | **ID** | **P.Value** |
| --- | --- | --- |
| **hsa-mir-152** | **hsa-miR-152_st** | **0.00000135** |
| **hsa-mir-708** | **hsa-miR-708_st** | **0.00020837** |
| **hsa-mir-21** | **hsa-miR-21-star_st** | **0.00021884** |
| **hsa-mir-200a** | **hsa-miR-200a-star_st** | **0.00026783** |
| **hsa-mir-339** | **hsa-miR-339-3p_st** | **0.00046164** |
| **hsa-mir-502** | **hsa-miR-502-3p_st** | **0.00054615** |
| **hsa-mir-151** | **hsa-miR-151-3p_st** | **0.00063115** |
| **hsa-mir-151** | **hsa-miR-151-5p_st** | **0.00105479** |
| **hsa-mir-138-1 // hsa-mir-138-2** | **hsa-miR-138_st** | **0.00108605** |
| **hsa-mir-151b** | **hsa-miR-151b_st** | **0.00115726** |
| **hsa-mir-3978** | **hsa-miR-3978_st** | **0.00189054** |
| **hsa-mir-452** | **hsa-miR-452_st** | **0.00195467** |
| **hsa-mir-130a** | **hsa-miR-130a_st** | **0.00217272** |
| **hsa-mir-3659** | **hsa-miR-3659_st** | **0.00231807** |
| **hsa-mir-135b** | **hsa-miR-135b_st** | **0.00238409** |
| **hsa-mir-4658** | **hsa-miR-4658_st** | **0.00281224** |
| **hsa-mir-500a** | **hsa-miR-500a_st** | **0.00302218** |
| **hsa-mir-183** | **hsa-miR-183_st** | **0.00334308** |
| **hsa-mir-34a** | **hsa-miR-34a_st** | **0.00336931** |
| **hsa-mir-1255b-1 // hsa-mir-1255b-2** | **hsa-miR-1255b_st** | **0.00348079** |
| **hsa-mir-4642** | **hsa-miR-4642_st** | **0.0036564** |
| **hsa-mir-500a** | **hsa-miR-500a-star_st** | **0.00388269** |
| **hsa-mir-182** | **hsa-miR-182_st** | **0.00393783** |
| **hsa-mir-362** | **hsa-miR-362-5p_st** | **0.00458656** |
| **hsa-mir-636** | **hsa-miR-636_st** | **0.0046749** |
| **hsa-mir-34a** | **hsa-miR-34a-star_st** | **0.00478709** |
| **hsa-mir-454** | **hsa-miR-454-star_st** | **0.00487284** |
| **hsa-mir-518f** | **hsa-miR-518f-star_st** | **0.00516263** |
| **hsa-let-7e** | **hsa-let-7e_st** | **0.00593499** |
| **hsa-mir-4266** | **hsa-miR-4266_st** | **0.00596467** |
| **hsa-mir-501** | **hsa-miR-501-3p_st** | **0.00715222** |
| **hsa-mir-10b** | **hsa-miR-10b_st** | **0.0074664** |
| **hsa-mir-4636** | **hsa-miR-4636_st** | **0.00778354** |
| **hsa-mir-516b-1 // hsa-mir-516b-2** | **hsa-miR-516b-star_st** | **0.00868822** |
| **hsa-mir-4523** | **hsa-miR-4523_st** | **0.00886037** |
| **hsa-mir-2682** | **hsa-miR-2682-star_st** | **0.00902375** |
| **hsa-mir-224** | **hsa-miR-224-star_st** | **0.00921168** |
| **hsa-mir-1301** | **hsa-miR-1301_st** | **0.00930066** |
| **hsa-mir-323b** | **hsa-miR-323b-3p_st** | **0.00939191** |
| **hsa-mir-4255** | **hsa-miR-4255_st** | **0.00088984** |
