## Supplementary material for "A miRNAs Based Exploration of promising Biomarkers in Cervical Cancer using Bioinformatic Methods": Table 2

| miRNA list | Area | Std.Error | 95% confidence interval | P value |
| --- | --- | --- | --- | --- |
| hsa_mir_152 | 1 | 0 | 1.000 to 1.0000 | 0.0126 |
| hsa_mir_21 | 1 | 0 | 1.000 to 1.0000 | 0.0126 |
| hsa_mir_339 | 1 | 0 | 1.000 to 1.0000 | 0.0126 |
| hsa_mir_151 | 1 | 0 | 1.000 to 1.0000 | 0.0126 |
| hsa_mir_138-1/138-2 | 1 | 0 | 1.000 to 1.0000 | 0.0126 |
| hsa_mir_3978 | 1 | 0 | 1.000 to 1.0000 | 0.0126 |
| hsa_mir_3659 | 1 | 0 | 1.000 to 1.0000 | 0.0126 |
| hsa_mir_135b | 1 | 0 | 1.000 to 1.0000 | 0.0126 |
| hsa_mir_4658 | 1 | 0 | 1.000 to 1.0000 | 0.0126 |
| hsa-mir-4266 | 1 | 0 | 1.000 to 1.000 | 0.0126 |
| hsa-mir-516b-1 // hsa-mir-516b-2 | 1 | 0 | 1.000 to 1.000 | 0.0126 |
| hsa-mir-2682 | 1 | 0 | 1.000 to 1.000 | 0.0126 |
| hsa-mir-518f | 1 | 0 | 1.000 to 1.000 | 0.0126 |
